# Nanoliposomal Omega-3 Fatty Acids Promote Adult Hippocampal Neurogenesis through the BDNF/TrkB Pathway in C57BL/6 Mice

**DOI:** 10.64898/2026.02.24.707750

**Authors:** RB Foltran, G Diaz, KM Stefani, MS Feliu, AR Impa Condori, AA Colapietro, DR Montagna, V Ambrosi, MF Godoy, SM Guidi, M Nanni, SL Diaz

**Affiliations:** Instituto de Fisiología, Biología Molecular y Neurociencias (IFIBYNE), UBA-CONICET, Buenos Aires, Argentina; Instituto Tecnología de Alimentos, INTA, Buenos Aires, Argentina; Instituto de Ciencia y Tecnología de Sistemas Alimentarios Sustentables (ICyTeSAS) INTA – CONICET, Buenos Aires, Argentina; Facultad de Farmacia y Bioquímica (FFyB), Cátedra de Bromatología, UBA. Buenos Aires, Argentina; Instituto de Biología Celular y Neurociencias (IBCN), UBA-CONICET, Buenos Aires, Argentina; Universidad Argentina de la Empresa (UADE). Buenos Aires, Argentina; Facultad de Farmacia y Bioquímica (FFyB), Cátedra de Nutrición, UBA. Buenos Aires, Argentina; Instituto de Biociencias, Biotecnología y Biología traslacional (IB3), UBA-CONICET. Buenos Aires, Argentina; Facultad de Ciencias Veterinarias (FVET), Cátedra de Química Biológica, UBA. Buenos Aires, Argetina; Universidad de Moron, Escuela Superior de Ingeniería, Informática y Ciencias Agroalimentarias (ESIICA, UM), Argentina; Universidad Nacional de Hurlingham (UNAHUR), Argentina; Facultad de Farmacia y Bioquímica (FFyB), Cátedra de Técnicas de Bioterio, UBA. Buenos Aires, Argentina

**Author notes:** These authors contributed equally to this article (RB Foltran, G Diaz).

## Abstract

Polyunsaturated fatty acids (PUFAs) are fundamental for different cellular and structural processes, especially regarding the nervous system. However, its incorporation in food has many bioaccessibility limitations, making it important to find new ways of its consumption. In this work, nanoencapsulated PUFAs orally administrated to C57BL/6 elite male mice for 8 weeks showed better bioavailability when compared to administration of free acids, also improving the effects on dentate gyrus neuronal survival as well as on the proneurogenic elements of the Brain derived neurotrophic factor biological pathway also in the hippocampus. In addition, nanoencapsulated PUFAs increased expression of Fabp5, a relevant n-3 fatty acids transporter in the brain. Altogether, our results would mean that the form of administration of the fatty acids can alter not only how much and how preserved they reach the central nervous system, but also have a differential impact in the diverse processes they contribute to.

## Introduction

Polyunsaturated fatty acids (PUFAs) are indispensable structural and signaling components of cellular architecture. Scientific interest in neurobiology focuses on omega-3 Polyunsaturated fatty acids (*n*-3 PUFAs), specifically, eicosapentaenoic acid (20:5, *n*-3; EPA) and docosahexaenoic acid (22:6, *n*-3; DHA). These molecules are far more than energy substrates: they are preferentially esterified into neuronal membrane phospholipids. Within this lipid bilayer, their high degree of unsaturation modulates critical physicochemical properties, i.e. membrane fluidity, curvature, and the spatial organization of lipid rafts, essential for neurotransmission and synaptic plasticity (1,2).

The physiological efficacy of n-3 PUFAs is dictated by their transition from the diet to the brain parenchyma, a process restricted by the blood-brain barrier (BBB). Recent evidence identifies the Major Facilitator Superfamily Domain-containing protein 2a (MFSD2A) as the primary transporter for lysophosphatidylcholine-bound DHA, while the Fatty Acid-Binding Protein 5 (*Fabp5*) facilitates the uptake of non-esterified forms (3,4). Collectively, this coordinated transport mechanism represents the physiological bottleneck that determines the neuroprotective efficacy of lipids.

Furthermore, the modern “Western Diet” has shifted the systemic lipid profile toward a pro-inflammatory state, characterized by an excessive *n*-6/*n*-3 ratio and high Saturated Fatty-Acids (SFA) intake, which respectively competes with *n*-3 PUFAs for desaturation and elongation pathways, and promote membrane lipid remodeling that limits n-3 PUFA incorporation (5). Conventional n-3 incorporation in food, however, is hindered by the extreme susceptibility of methylene-interrupted double bonds to oxidative stress. The resulting lipid peroxidation products not only diminish biological functionality but can also induce cellular toxicity.

To overcome these bioaccessibility limitations, the use of fermented dairy matrices offers a promising strategy as a carrier for omega-3 fatty acids, owing to their ability to optimize bioactive stability and functionality. Furthermore, nanotechnology-specifically through nanoemulsions and encapsulation-has been established as a superior strategy to enhance lipid delivery (6). By protecting PUFAs from gastric degradation and optimizing intestinal micellization, these systems significantly increase the concentration of active fatty acids in target tissues (7).

A central objective of this research is to elucidate how this enhanced omega-3 fatty acids delivery impacts adult hippocampal neurogenesis in a murine model. The dentate gyrus (DG) maintains a neurogenic niche where multipotent stem cells expressing Nestin (Nes) transition into mature, Calbindin-positive (Calb1) functional neurons (8). This continuum is orchestrated by the Brain-Derived Neurotrophic Factor (BDNF) pathway, among other factors. The BDNF is synthesized as a pro-protein, the 32 kDa pro-BDNF that may be cleaved to yield the 13.2-15.9 kDa mature BDNF (mBDNF). The pro-BDNF can release mBDNF after cleavage either intra- or extra-cellularly, or be secreted as pro-BDNF without further modifications. By binding to TrkB receptors, the mBDNF facilitates pro-neurogenic effects, whereas the pro-BDNF may bind different receptors, being the most known the p75, through which pro-apoptotic actions are promoted (see a rev. in (9)). While DHA is known to upregulate BDNF (10,11), the specific molecular mechanisms by which nanoencapsulated n-3 PUFAs (DHA-EPA) modulate the expression of markers of neuronal differentiation during this process remain under-investigated. Therefore, the present study evaluates the effects of orally administered nanoencapsulated n-3 PUFAs (DHA-EPA), delivered via a fermented yogurt matrix, in male C57BL/6 mice over a chronic period. The administration of the bioactive compounds was based on the Recommended Daily Intake (RDI) of at least 250mg/day of EPA+DHA for healthy adults (excluding non-pregnant/non-lactating adult females) established by international organizations (12), integrated into a fermented dairy matrix to ensure its bioavailability and mimic an optimized human consumption scenario. We hypothesize that this formulation increases systemic bioavailability, BBB transport efficiency, and metabolic incorporation compared to free fatty acids, thereby enhancing neuronal survival in the DG and robustly activating the BDNF signaling cascade.

## Materials and Methods

### Animals

Studies were performed on 51 male, C57BL/6 elite mice purchased at the Instituto de Medicina Experimental, Academia Nacional de Medicina, Buenos Aires, Argentina. Experiments on animals were conducted according to local regulations and were approved by the Institutional Ethical Committee (ResCD-2022-1246-E-UBA-DCT#FMED). Three-four-week-old mice, bred in barrier-conditions to maintain an SPF status, were transported in environmentally controlled conditions to our institute’s animal facility. After arrival, mice were housed in 1284L Eurostandard Type II Long (365 mm × 207 mm × 140 mm) Tecniplast microisolator cages with filter tops (five to six animals per cage), with autoclavated aspen shavings as bedding and tissue paper as nesting material. Mice were maintained under controlled conditions, i.e., 22 ± 2°C room temperature, 40-70% relative humidity, 12–12 h light–dark cycle (lights on at 8 a.m.), pelleted food for rodents (Cooperación) and water ad libitum. Cages were changed twice a week. A period of acclimation of 2 weeks was left before the beginning of experiments, and therefore, mice were 5–6 week-old when treatments began.

### Yogurt preparation and administration

#### Raw Materials and Starter Culture

Yogurt was manufactured using commercial pasteurized cow milk (2% w/v fat; *Mastellone Hnos. S.A., Buenos Aires, Argentina*). A lyophilized Direct Vat Set (DVS) thermophilic starter culture, YoFlex® YG-X16 (comprising *Streptococcus thermophilus* and *Lactobacillus delbrueckii* subsp. *bulgaricus*), was provided by Chr. Hansen (Hørsholm, Denmark). Prior to inoculation, the culture was activated by hydrating 3 g of the lyophilized powder in 1000 mL of milk at 45 °C according to the manufacturer’s instructions.

#### Functional Ingredients and Yogurt Preparation

Omega-3 oil (MEG-3™ 2050EE) was sourced from DSM Nutritional Products. Three distinct formulations were established based on the omega-3 delivery system:

- Control: Yogurt without oil supplementation.
- Free Acids: Yogurt supplemented with bulk omega-3 oil (250 mg / 200 mL milk).
- Nano Acids: Yogurt supplemented with a nanoencapsulated omega-3/soy lecithin formulation. To achieve an equivalent dose of 250 mg of active omega-3 per 200 mL (provided by Nanotica Agro SRL via an R&D agreement with INTA) was incorporated.

In both functional formulations, bioactive compounds were homogenized into the milk-sugar base (75 g/L) prior to fermentation.

### Yogurt Manufacturing

The mixtures were inoculated with the activated starter culture (2 mL/L) and aseptically distributed into 200 g sterile polypropylene containers. Incubation was performed at 43 ± 1 °C until a target pH of 4.5 was achieved. Samples were then cooled and stored at 4 °C for 28 days for analysis.

### Yogurt Administration

The different formulations of yogurt were administered to mice by mixing it with a palatable cube. They were made with 10 g of powdered yeast, 66 g of powdered milk, 36 g of sugar, and 14 g of gelatin, for each ice tray (12 cubes) (13). These ingredients were mixed with 2 ml of each of the three yogurts and with approximately 60 mL of tap water, forming a thick liquid preparation, which was poured into an ice tray. The 3 cm × 3 cm × 4 cm cubes, weighing around 12 g, were kept at 4 °C and were prepared every 6 days. A yogurt containing cube was given per cage, every day between 2 p.m. and 5 p.m. Mice voluntarily ate the cubes, immediately after they were placed in the cages.

### Experiments

All mice were divided into 3 groups (control vs the two types of yogurt formulation) for 3 different experiments. For the first protocol, 5 mice per group were administered the yogurt cubes for 4 weeks. At the end of the treatment, all mice were deeply anesthetized (20 mg/kg xylazine and 200 mg/kg ketamine) and blood was obtained by intracardiac collection, from which serum was extracted to quantify fatty acids concentrations. After death confirmation, mice were decapitated, brains were removed, and both hippocampus (HC) were obtained, where the left hemi-HCs were prepared for protein extraction and Western blots and the right hemi-HCs were used for mRNA measurements by qRT-PCR.

For the second experiment, in order to study neuronal proliferation 18 mice (n = 6 per group) were administered yogurt cubes for 4 weeks. The thymidine analog 5-bromo-2′-deoxyuridine (BrdU; Sigma, B9285) was administered to mice at the end of the treatment, in order to label dividing cells. Animals received two ip injections of BrdU, 50 mg/kg each, at a 2 h interval. BrdU was dissolved in 0.9% NaCl at 50 °C. The next day, all mice were deeply anesthetized (20 mg/kg xylazine and 200 mg/kg ketamine) and transcardially perfused with 5 mL of 0.9% NaCl with heparin and 50 mL of 4% paraformaldehyde in 0.1 M phosphate-buffered saline (PBS, pH 7.4) for 15 min. Brains were recovered, post-fixed for 24 h at 4 °C, and cryoprotected with a 30% sucrose solution. Brains were then sliced into 35 μm thick coronal sections through the hippocampus (bregma −0.94 to −3.80) with a freezing microtome. Sections were stored at 4 °C in 0.1 M PBS with 0.1% sodium azide if not used immediately.

For the last trial, the yogurt cubes were administered for 8 weeks to 18 mice (6 per group) to study the survival of newborn neurons. Animals received two ip injections of BrdU, 50 mg/kg each, at a 2 h interval for two consecutive days at the end of the 4th week of treatment. Mice were then continued treatment for 4 more weeks, so that BrdU labeled 4 week-old neurons. At the end of the treatment, all mice were deeply anesthetized and transcardially perfused, as stated before.

### Fatty acid profile determination

Serum fatty acid profile was determined by gas chromatography: Chromatograph: Perkin Elmer Clarus 500; Column: Supelco SP 2560 100 m x 0.25 mm x 0.20 μm; FID detector 280 °C; Carrier gas: helium. Previously, serum lipids were converted into methyl esters according to the Lepage and Roy method as follows: 2 mL of methanol/toluene (4:1) were added to 200 µL of serum, followed by 0.2 mL of acetyl chloride using a vortex mixer. The mixture was incubated at 100 °C for one hour. Then, 5 mL of 6% potassium carbonate was added, and the samples were centrifuged; 1 µL of the toluene layer was injected into the chromatograph. Fatty acid methyl esters (FAMEs) were identified based on their retention times by comparison with the PUFA No. 2 (Animal Source) FAME analytical standard (SUPELCO 47015-U). The results (mean ±SD) were expressed as the per cent (%) of total fatty acids content, with a quantification limit of 0.05% (14).

### Western Blotting

Tissue was homogenized with 250 μl of RIPA buffer (150 mM NaCl, 1% NP-40, 0.5% Sodium deoxycholate, 0.1% SDS, 50 mM Tris) along with protease inhibitors (1:100) and centrifuged at 4°C for 30 min at 13000 (r/min). The supernatants were recovered and protein levels were quantified by the Bradford protein assay. Samples (50 μgr in 5× loading buffer) were then loaded into SDS-PAGE gels (12 or 15%) and transferred onto nitrocellulose membranes using the Mini-PROTEAN® Tetra System (BIO-RAD) for 1 h.

Membranes were incubated for 1 h with blocking solution (5% milk in TBST) and then probed overnight at 4°C with mouse anti-BDNF (1:2000; Icosagen; 327-100 clone 3C11), rabbit anti-p75 (1:700; Alomone Labs; ANT-007), rabbit anti-TrkB (1:700; Alomone Labs; ANT-019), and rabbit anti-proBDNF (1:2500; Genocopoeia) in TBST. Actin was used as a loading control (1:2000; Santa Cruz). Binding of primary antibodies was visualized with anti-mouse HRP-conjugated secondary antibody (1:20000; Invitrogen) or anti-rabbit HRP-conjugated secondary antibody (1:20000; Invitrogen). Membranes were developed using the ECL Plus Western blotting substrate (Thermo Fisher Scientific) for chemiluminescence with the Gene Gnome® Imager. Densitometry was carried out using ImageJ software (15). The signal of each protein is expressed after subtraction of background signal and related to actin signal.

### qRT-PCR

Hippocampal tissue was collected in 250 µL cold Trizol (Invitrogen, Paisley, UK) and homogenized using plastic pestles for homogenizers. RNA was isolated according to the manufacturer’s protocol. After addition of chloroform, samples were centrifuged at 11,000 × g for 15 min at 4 °C. The aqueous phase was transferred and RNA was precipitated with isopropanol, washed with 75% ethanol and resuspended in 15 µL of RNase-free DEPC-treated water. RNA concentration of the tissue was quantified with the Qubit 2.0 Fluorometer (RNA Assay BR Kit, Invitrogen). To ensure genomic DNA removal, 1 μg of total RNA was treated with 1 U/μL of DNase I (Invitrogen) for 15 min. RNA purity and absence of residual DNA were verified by qPCR using non-reverse-transcribed RNA as a negative control. Samples were stored at −80 °C until further use.

Reverse Transcription Quantitative Polymerase Chain Reaction (qRT-PCR) was carried out in a StepOnePlus Real-Time PCR System (Applied Biosystems, CA, USA), and the data were analyzed with StepOne software v2.2. The retrotranscription and amplification reactions were performed in a total volume of 10 μl buffer solution containing 2 μl of template RNA, 5 μl of 2× iTaq universal SYBR® Green reaction mix (2x), 0.125 μl iScript reverse transcriptase (Applied Biosystems), 2.08 μl RNase-free DEPC-treated water and 0.4 μl of each primer (400 μM). Primers for BDNF (Brain-Derived Neurotrophic Factor), FABP5 (Fatty Acid Binding Protein 5), Calb1 (Calbindin) and Nes (Nestin) were designed using Primer Express software (Applied Biosystems) and described in table 1.

**Table 1.**
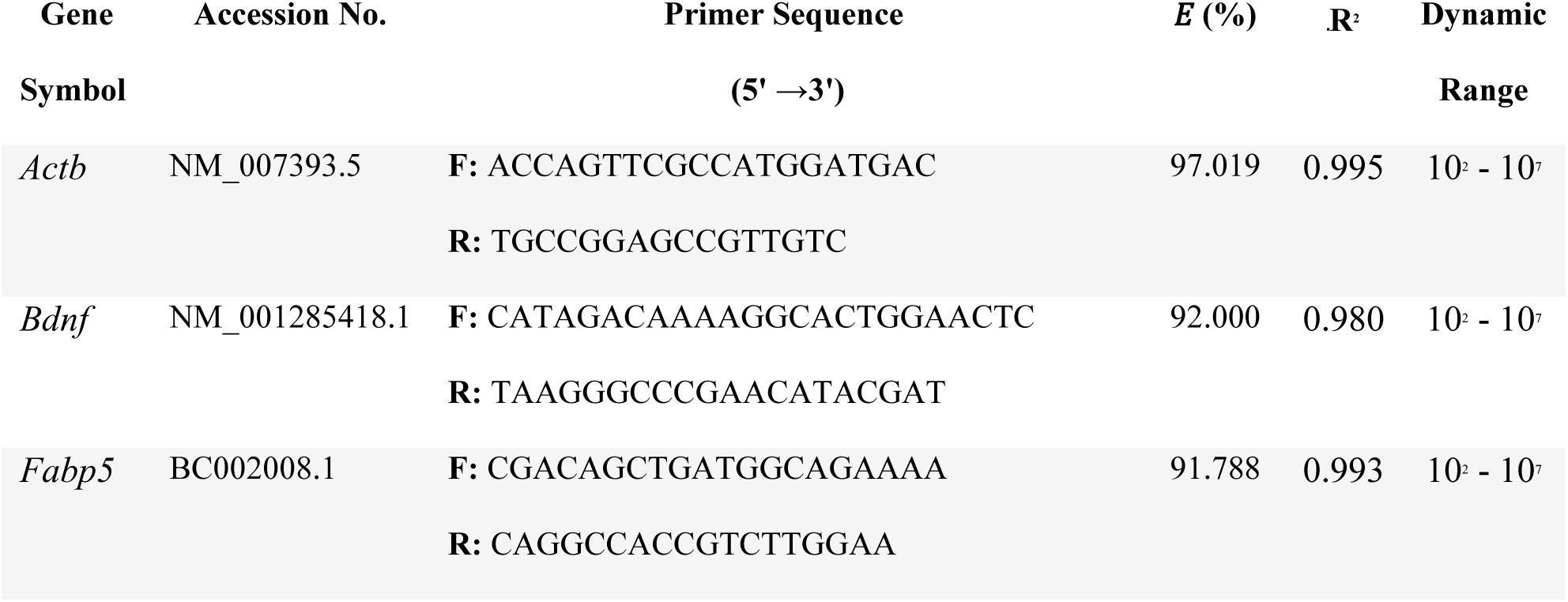
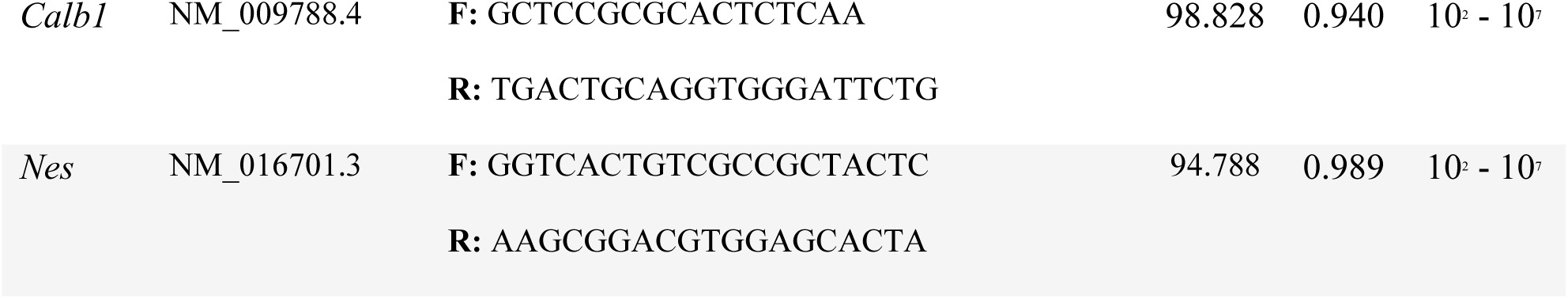
Primer sequences and qPCR performance characteristics. **Accession No.** refers to the NCBI GenBank database. ***E* (%)** represents the amplification efficiency calculated from the slope of the standard curve (E=[10^(-1/slope)^−1]×100. ***R* ^2^** denotes the coefficient of determination for the linear regression. **Dynamic Range** indicates the interval of template concentration where linearity was maintained. All assays were performed in technical triplicate using SYBR Green chemistry on a StepOnePlus™ Real-Time PCR System.

The thermal cycling profile consisted of:

- UNG pre-incubation: 50 °C for 2 min.
- Initial denaturation: 95 °C for 1 min.
- Amplification (40 cycles): 95 °C for 15 s and 60 °C for 1 min.
- Melt curve analysis: 95 °C (15 s), 60 °C (1 min), and a final ramp to 95 °C at 0.6 °C/s to confirm amplicon specificity.

To ensure the robustness of the gene expression analysis, three independent biological replicates were processed per experimental group, following the recommendations of the MIQE guidelines (16). Each biological sample was analyzed in triplicate using qRT-PCR. This balanced design allows for an accurate estimation of experimental error even with biological sample sizes of n=3, ensuring that the observed differences possess true statistical and biological significance. Target gene expression was normalized against *β*-actin (ACTB) as the internal reference gene. The dynamic range was established using a six-point standard curve derived from three independent biological replicates, each processed in technical triplicate (n=9 per dilution). R^2^ values and amplification efficiencies were calculated based on mean Cq values, incorporating a housekeeping gene (also analyzed in triplicate) for normalization. CT (threshold cycle) values were calculated by subtracting the CT value of ACTB from the CT values. Relative expression was calculated as E^−ΔCt, where E is the efficiency measured for each set of primers.

### Immunofluorescence

Newborn cells in the dentate gyrus (DG) were revealed by immunostaining against BrdU as previously published (17). Free-floating sections were exposed to 2 N HCl for 1 h. Nonspecific binding was reduced by 1 h incubation in a saturation solution containing 2 g/L gelatin (Merck) and 0.25% triton in PBS. Sections were then incubated overnight at 4 °C with a mouse anti-BrdU antibody (1:1000; Developmental Studies Hybridoma Bank, concentrated). Next day, the tissue was incubated for 2 h with an anti-mouse-A488 (1: 500; Invitrogen). All primary and secondary antibodies were diluted in the saturation solution. Finally, sections were incubated with Hoechst (1: 10000, Invitrogen) to dye the nucleus of cells. The number of BrdU-labeled cells was quantified with a fluorescence microscope (Olympus IX81) with a 20× objective, on serial sections through the entire hippocampus. One out of six series was counted (180 μm intervals, 10 sections per series). BrdU-labeled cells were counted in the sub-granular zone of the granule-cell layer over the entire DG. Cells were considered to be BrdU^+^ when their nuclei were completely filled with fluorescence or showed clear patches of variable intensity (classic dilution pattern for survival assays).

### Statistical Analysis

Group differences were analyzed using one-way analysis of variance (ANOVA). Assumptions of normality and homogeneity of variances were assessed using the Shapiro–Wilk and Bartlett tests, respectively. When variance heterogeneity was detected, statistical analyses were performed using either Welch’s one-way ANOVA or generalized least squares (GLS) models that explicitly accounted for heteroscedasticity. For GLS analyses, models allowing different variance structures were compared to homoscedastic models using likelihood ratio tests, and the final model was selected by balancing goodness-of-fit and parsimony. Post hoc comparisons were performed using Tukey’s test following classical ANOVA and GLS models, whereas Games–Howell tests were used following Welch’s ANOVA. In fatty acid analyses, p-values were adjusted for multiple testing using the Benjamini–Hochberg false discovery rate (FDR) procedure, applied separately to individual fatty acids and fatty acid classes. Body weight was analyzed using a two-way repeated-measures ANOVA with week as a within-subject factor and treatment as a between-subject factor. Sphericity was assessed using Mauchly’s test and Geenhouse-Geisser corrections were applied when necessary. In all analyses, p < 0.05 was considered statistically significant, whereas values between 0.05 and 0.1 were considered a trend.

## Results

### Body weight

Mice were weighed across every week of treatment to control for potential changes due to the different treatments. Analyses showed an interaction between treatment and number of weeks (p = 0.030), with no difference in treatment (p = 0.145) but significant changes across weeks (p <.001). As expected, given their age, all mice gained some weight along the experimental protocol but no differences were revealed between experimental groups.

### Fatty acids profile

Individual serum levels of the SFA lauric (C12:0), myristic (C14:0) and palmitic (C16:0) were not altered by the treatments (p = 0.2482, p = 0.5138, p = 0.5962), whereas the SFA stearic (C18:0) and lignoceric (C24:0) showed a trend to higher values in the group that consumed the nanoencapsulated version of omega-3 fatty acids in yogurt compared to the control group (p = 0.0697) (Table 2).

**Table 2:** Fatty acid profile in serum of mice treated with different formulations of yogurt. Results are expressed as the percentage (%) of total fatty acids content. Data are expressed as mean ± SD, n =4-5/group. # 0.05 < p < 0.10; *p < 0.05; **p < 0.01; ***p < 0.001.

| Serum composition | Control | Free fatty acids | Nano fatty acids | Statistics | P value (FDR) |
| --- | --- | --- | --- | --- | --- |
| Lauric acid | 0.07 ± 0.023 | 0.072 ± 0.026 | 0.052 ± 0.004 | $F_2 = 2.538$ | 0.2482 |
| Myristic acid | 0.316 ± 0.056 | 0.355 ± 0.066 | 0.304 ± 0.036 | $F_2 = 1.1$ | 0.5138 |
| Palmitic acid | 18.2 ± 0.765 | 19.2 ± 1.39 | 19 ± 1.72 | $F_2 = 0.713$ | 0.5962 |
| Stearic acid | 8.48 ± 0.287 | 9.61 ± 0.972 | 9.46 ± 0.446* | $F_2 = 9.025$ | 0.0697 |
| Lignoceric acid | 0.166 ± 0.044 | 0.205 ± 0.031 | 0.284 ± 0.078* | $F_2 = 5.692$ | 0.0697 |
| Oleic acid | 17.3 ± 1.01 | 14.7 ± 1.94<sup>#</sup> | 14.6 ± 1.39* | $F_2 = 5.262$ | 0.0697 |
| Palmitoleic acid | 3.71 ± 0.430 | 3.66 ± 0.512 | 3.37 ± 0.414 | $F_2 = 0.786$ | 0.5962 |
| Linoleic acid | 20.9 ± 1.23 | 21.6 ± 1.97 | 21.5 ± 0.632 | $F_2 = 0.374$ | 0.7495 |
| Alfa-linolenic acid | 0.622 ± 0.108 | 0.598 ± 0.045 | 0.684 ± 0.084 | $F_2 = 1.24$ | 0.5138 |
| Arachidonic acid | 14 ± 1.72 | 12.3 ± 1 | 12.9 ± 2 | $F_2 = 1.166$ | 0.5138 |
| EPA | 0.514 ± 0.195 | 0.738 ± 0.033 | 0.75 ± 0.052 | $F_2 = 3.419$ | 0.1633 |
| DHA | 3.83 ± 0.243 | 4.2 ± 0.395 | 5.14 ± 0.262*** | $F_2 = 25.39$ | 0.00007 |

Both omega-3–fortified yogurt groups exhibited a trend to lower serum oleic acid (C18:1 n-9) levels compared with the control group (p = 0.0697), whereas no effect was observed on palmitoleic acid (C16:1 n-7) concentrations (p = 0.5962) (Table 2).

Regarding the essential PUFAs linoleic acid (C18:2 n-6) and α-linolenic acid (C18:3 n-3), no significant differences were observed among groups (p = 0.7495, p = 0.5138). Likewise, arachidonic acid (C20:4 n-6) levels were not significantly affected by the treatments (p = 0.5138) (Table 2).

Eicosapentaenoic acid (EPA, C20:5 n-3) showed a trend toward higher serum concentrations in the supplemented groups compared with the control (ANOVA p = 0.07; FDR-adjusted p = 0.1633) (Figure 1.A; Table 2). The control group exhibited intra-group variability, whereas both omega-3-enriched yogurt formulations resulted in more homogeneous and consistently elevated EPA levels. Although these differences did not reach statistical significance after multiple-comparison correction, the observed pattern suggests a potential treatment-related effect on circulating EPA.

**Figure 1.**
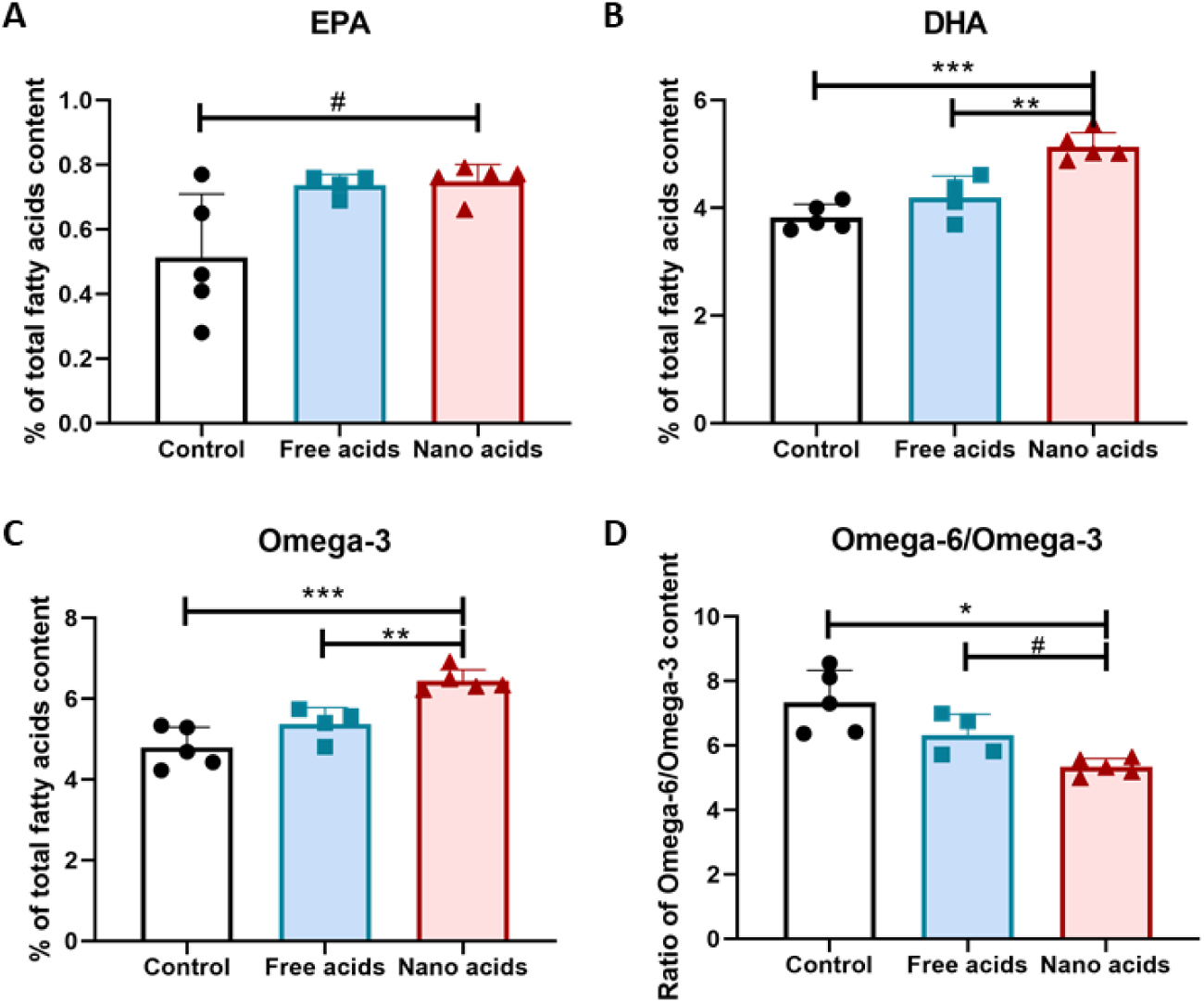
Omega-3 fatty acids profile and n-6/n-3 ratio in serum. Results are expressed as the percentage (%) of total fatty acids content. (A-B) Individual % for DHA and EPA. (C) Total % of fatty acids from the Omega-3 family. (D) Ratio of Omega-6/Omega-3 content in serum. Data are expressed as mean ± SD, n =4-5/group. # 0.05 < p < 0.10; *p < 0.05; **p < 0.01; ***p < 0.001

Docosahexaenoic acid DHA (C22:6 n-3) levels showed no significant differences in the free acids group compared to the control group (p = 0.1973) (Figure 1.B). However, the nanoencapsulated yogurt formulation produced a significant increase in serum DHA levels compared with both control and free-fatty acids groups (p = 0,00007). As expected, the group consuming the nanoencapsulated omega-3 yogurt exhibited a higher total n-3 fatty acid content (Figure 1.C) and a lower n-6/n-3 ratio (Figure 1.D), relative to the other groups. This finding suggests that nanoencapsulation may protect DHA from degradation during storage and digestion, thereby enhancing its stability and facilitating intestinal absorption, as reflected by increased serum DHA levels.

### Adult Hippocampal Neurogenesis

In order to analyze whether the fatty acid formulation differently altered the process of neurogenesis, BrdU^+^ cells were quantified by immunohistochemistry in the DG of mice, both at 24 h or 4 weeks after BrdU injection to respectively evaluate the proliferative and survival stages of the neurogenesis process. No significant changes were found in the proliferation of new cells (p = 0.67) but it should be noted that inspection of individual data points (Fig. 2A) revealed considerable within-group variability in all the experimental groups. Regarding survival of the newborn cells, we found a tendency to an increase in the mice treated with the nanoencapsulated formulation compared to control (ANOVA p = 0.0921, Post-hoc p = 0.0953941) (Fig. 2B) (Table 3).

**Figure 2:**
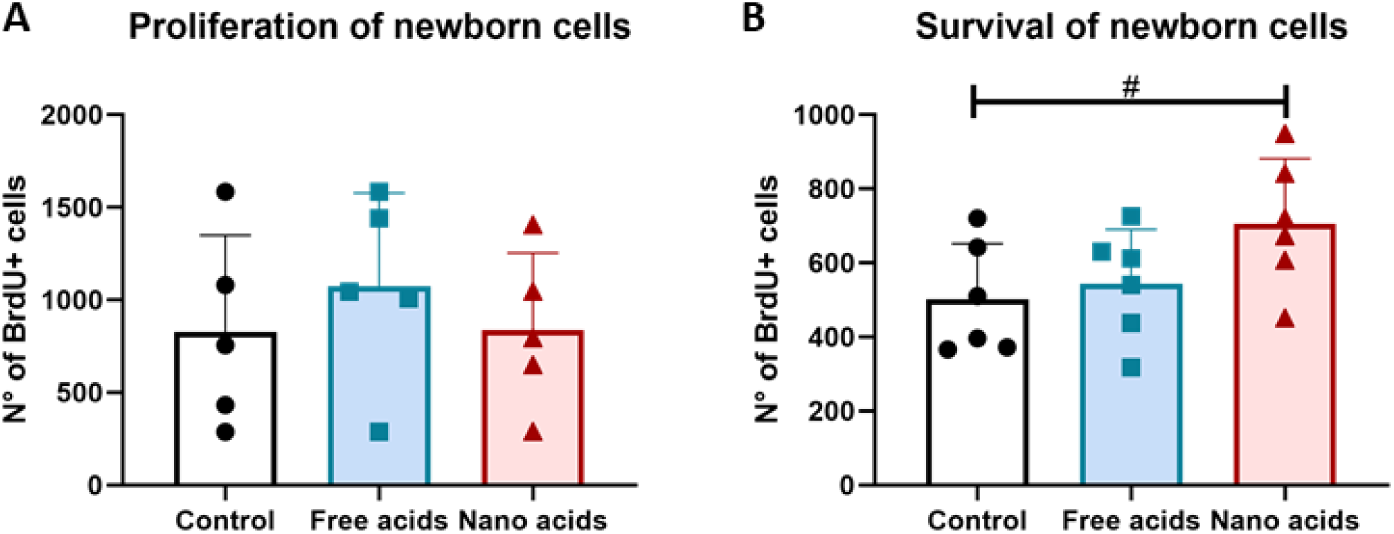
Adult neurogenesis process. (A) Proliferation of new cells in the dentate gyrus (DG) of mice treated with fatty acids supplementation, 24 hs post injection of BrdU. (B) Survival of newborn cells in the DG, 4 weeks post injection of BrdU. Data are expressed as mean ± SD, n =5−6/group. # 0.05 < p < 0.10.

**Table 3:**
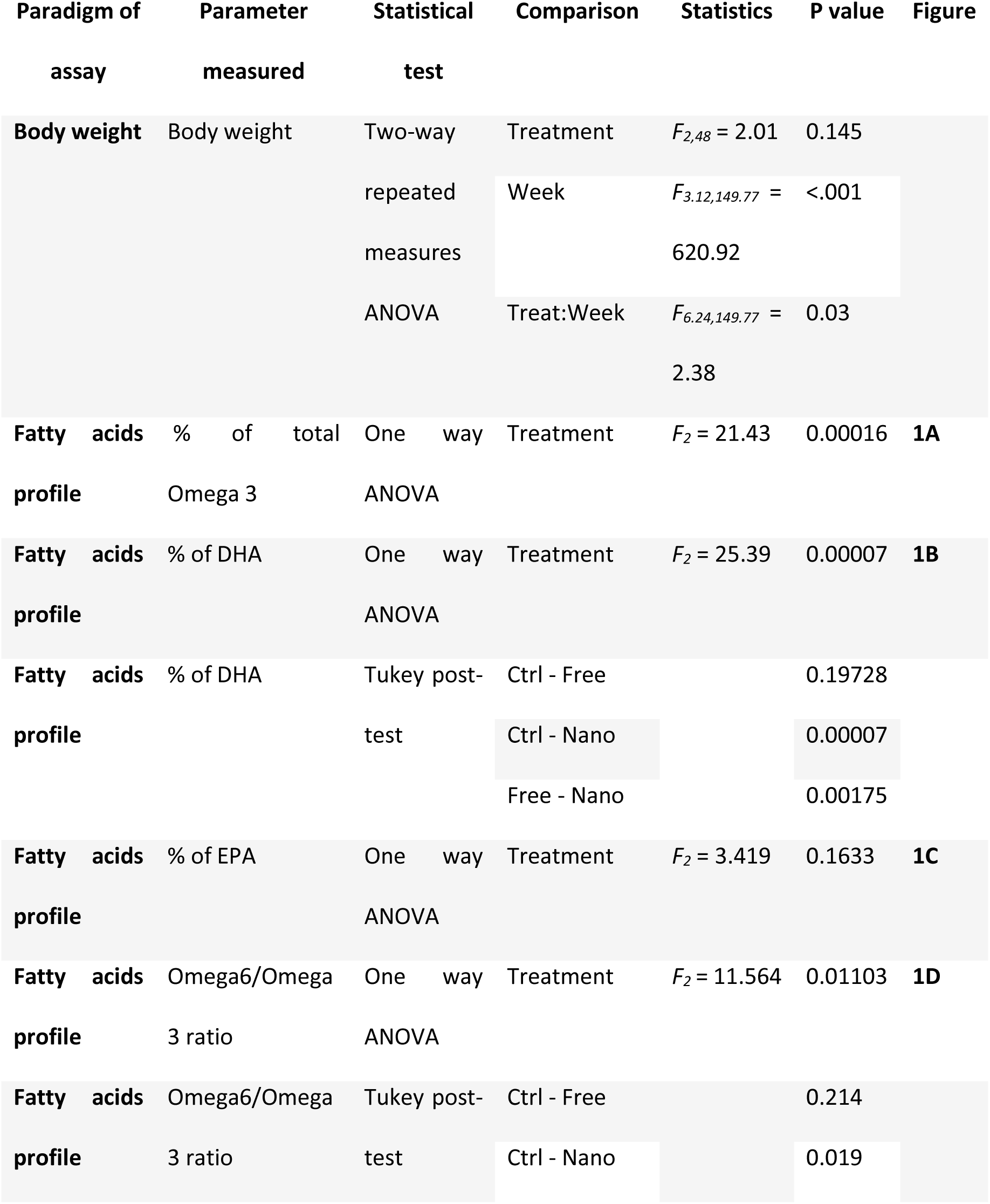
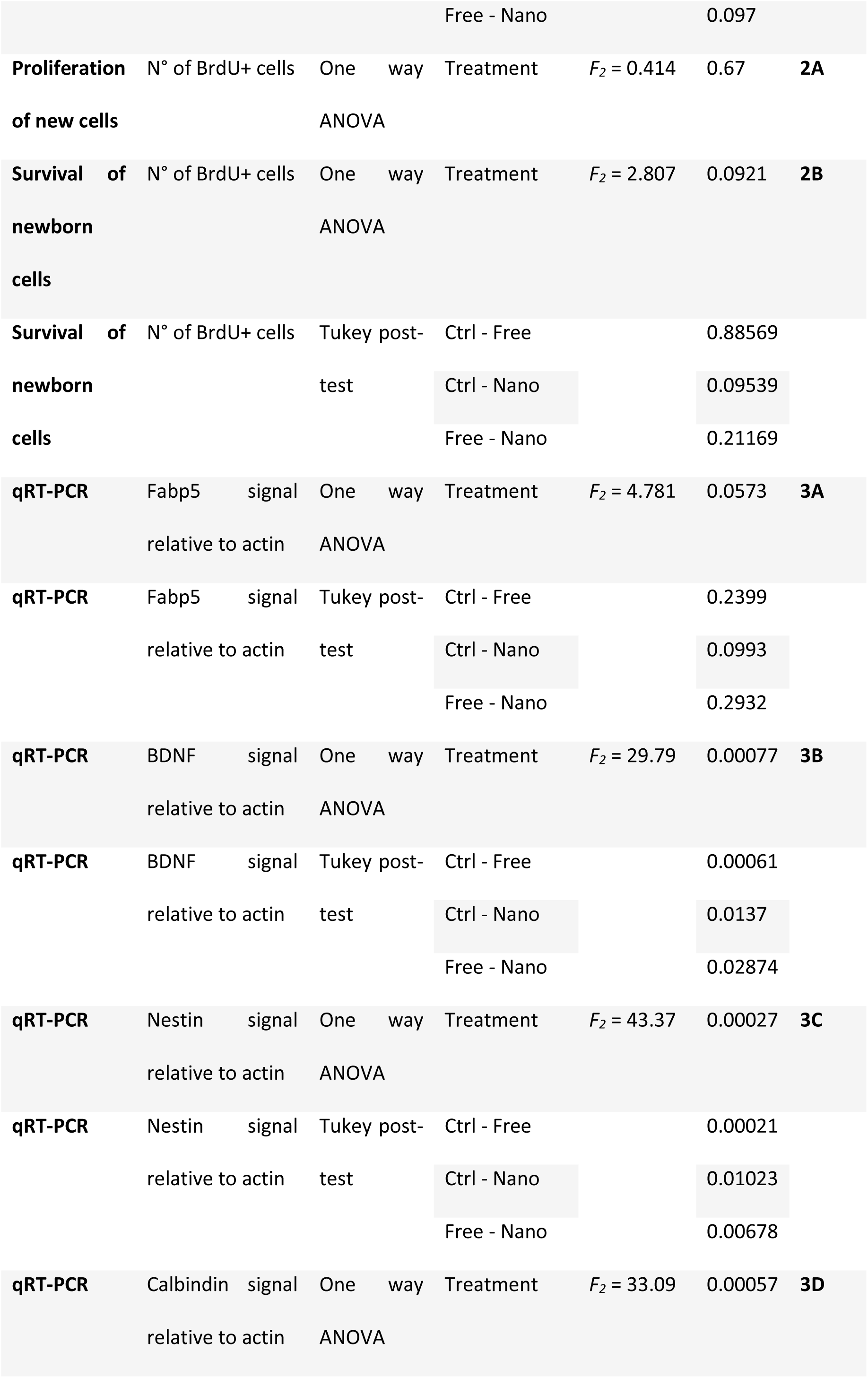
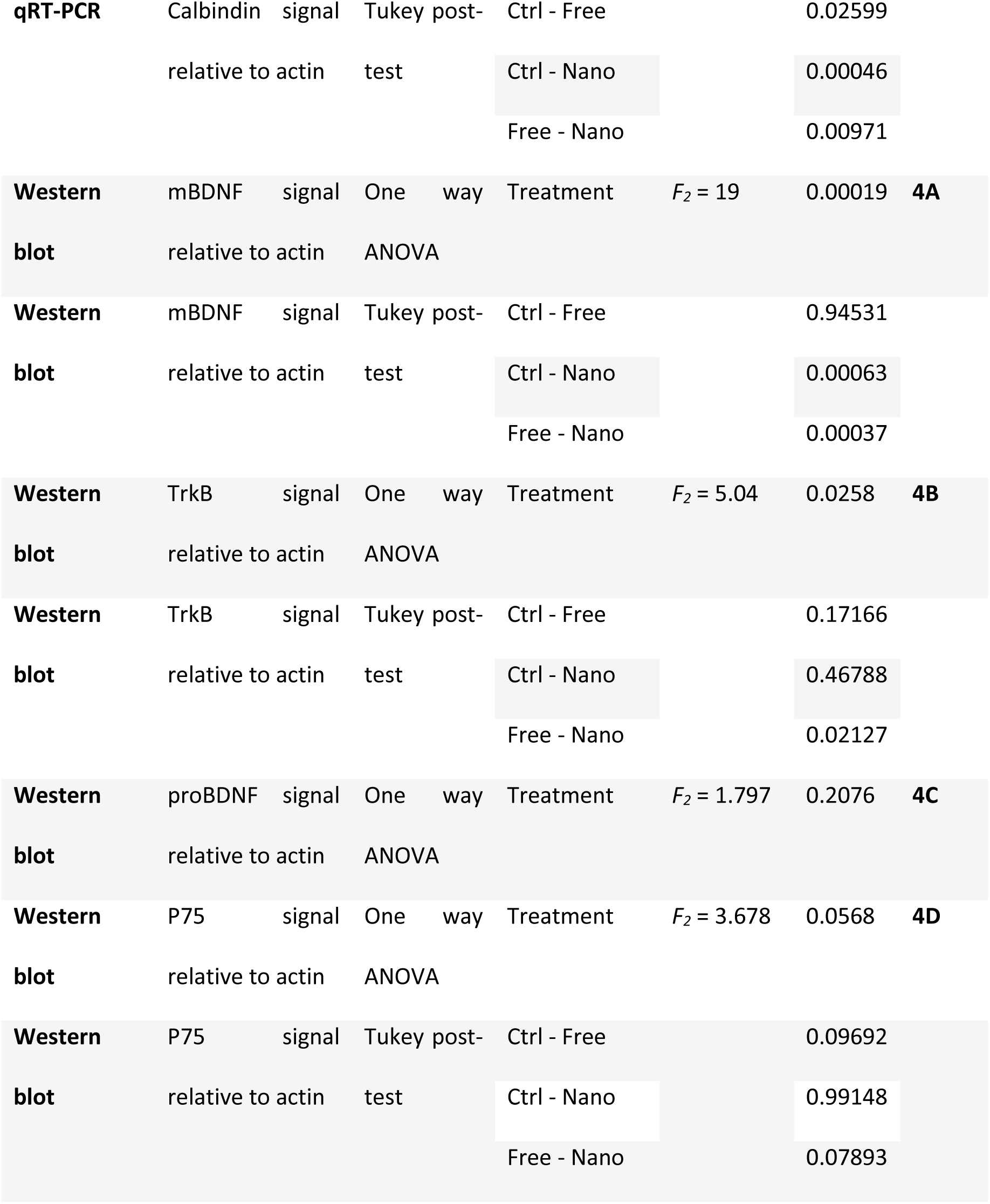
P values and statistical information of all procedures performed.

### RNA analysis

To have an insight into mechanisms behind fatty acid fortification effects in adult neurogenesis, the HC of mice were recovered after the 4 weeks of treatment and RNA expression of different key molecules involved in fatty acid transport, regulation of neurogenesis, and markers of young and mature neurons were analyzed.

As stated before, FABP5 plays a role in the uptake, transport, and metabolism of fatty acids, in particular, of DHA. A tendency to a higher FABP5 expression in mice treated with the nanoencapsulated yogurt formulation was detected (ANOVA p = 0.0573, Post hoc p = 0.0993) (Fig. 3A) (Table 3).

**Figure 3:**
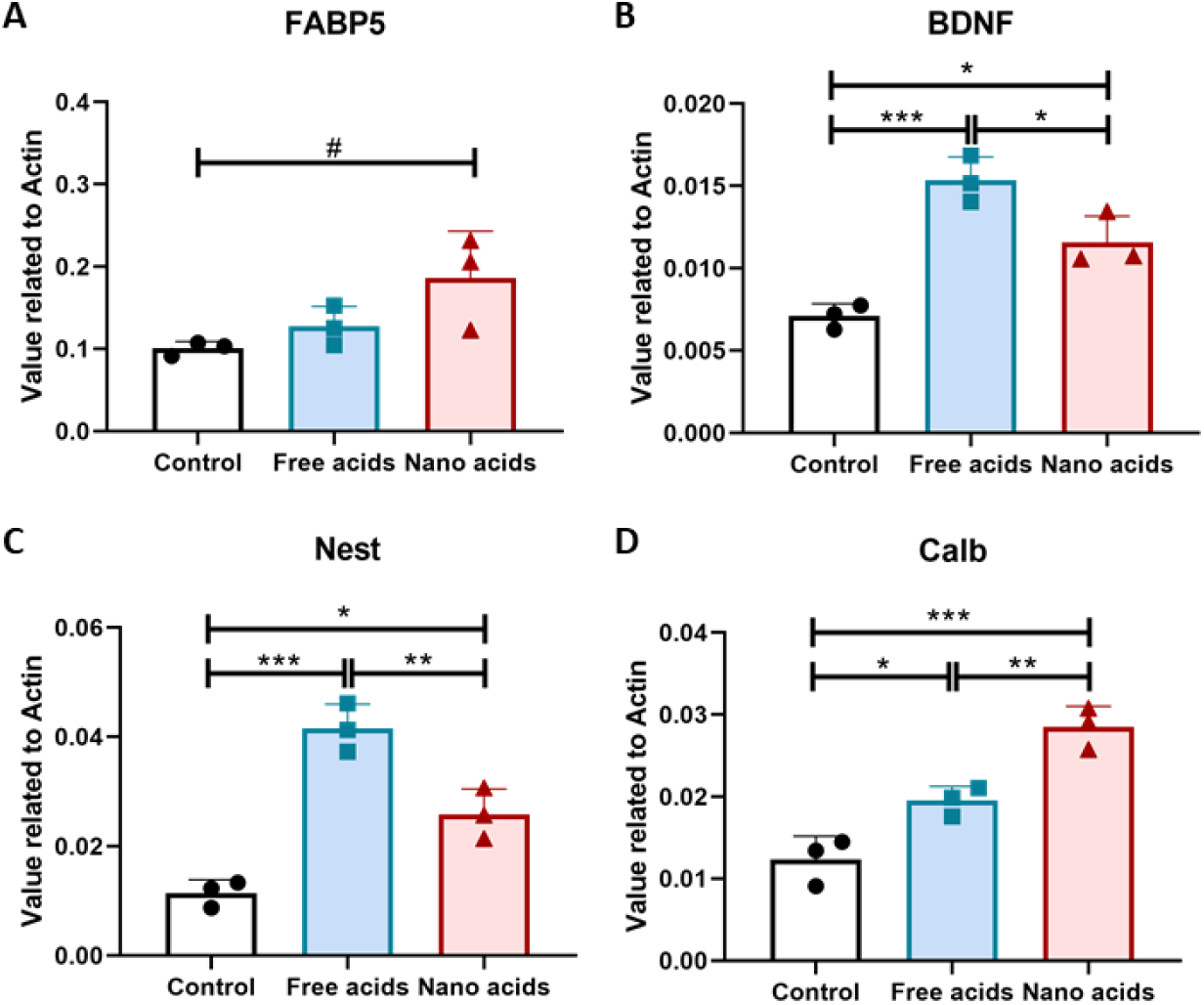
RTqPCRs. RNA was extracted from the HC of mice from the different groups to analyze the transcription levels of (A) FABP5, (B) BDNF, (C) Nestin and (D) Calbindin. Data are expressed as mean ± SD, n =3/group. # 0.05 < p < 0.10; *p < 0.05; **p < 0.01; ***p < 0.001

On the other hand, BDNF has been linked to the regulation of adult hippocampal neurogenesis which makes it a big candidate for being involved in the modulation of fatty acids effects in the brain. Indeed, we found a significant increase in BDNF RNA after the treatment (p = 0.000766) (Fig. 3B). Even though the neurotrophin expression levels were enhanced in both yogurt treated groups compared to control (Free-acids p = 0.0006111; Nano-acids p = 0.0137006), expression was higher after the free-fatty acids formulation (Free vs Nano p = 0.0287450) (Table 3).

Nestin is an intermediate filament protein that identifies neuronal progenitors and young newborn neurons, while Calbindin is a calcium binding protein which serves as a marker of mature, differentiated neurons. As a complementary approach to the immunohistochemistry analysis, and to study the molecular profile of the tissue, we measured RNA expression of both proteins. We found a significant increase in Nestin and Calbindin RNA levels after both fatty acid supplementation treatments (ANOVA Nes p = 0.000271, Calb p = 0.000575). For Nestin, the free-fatty acid formulation showed a higher expression (p = 0.0067782) (Fig. 3C) than the nanoencapsulated acids formulation, whereas Calbindin levels were more enhanced in the nanoencapsulated group (p = 0.0097060) (Fig. 3D) (Table 3). The increased expression of both neuronal markers are in line with the proneurogenic effect of EPA-DHA chronic administration revealed by immunohistochemistry.

### BDNF Pathway

To deepen the understanding of the changes produced in the BDNF pathway, which has a very complex regulation, protein-expression of both BDNF isoforms and their receptors was analyzed in the HC of yogurt-treated mice. Regarding the survival pathway, a significant increase was observed in mBDNF levels (ANOVA p = 0.000191) only in the animals treated with the nanoencapsulated yogurt formulation compared to controls (p = 0.0006301) (Fig. 4A, F), whereas changes in the TrkB receptor between the treated groups and the control mice were not significant (p = 0.4678826) (Fig. 4B, E) (Table 3). On the other hand, for the apoptotic pathway, no significant changes were found for proBDNF (p = 0.2076) though inspection of individual data points revealed considerable within-group variability in the free fatty acids group (Fig. 4C, F). In addition, a tendency towards a decrease in the p75 receptor was observed in mice treated with the free-fatty acid yogurt (ANOVA p = 0.0568, Ctrl vs Free p = 0.0969230) (Fig. 4D, F) (Table 3), which would mean less activation of the cell death mechanisms. Results show that the nanoencapsulated formulation promotes the modulation of the neurotrophin towards the survival pathway, which is in accordance to the previous outcomes.

**Figure 4:**
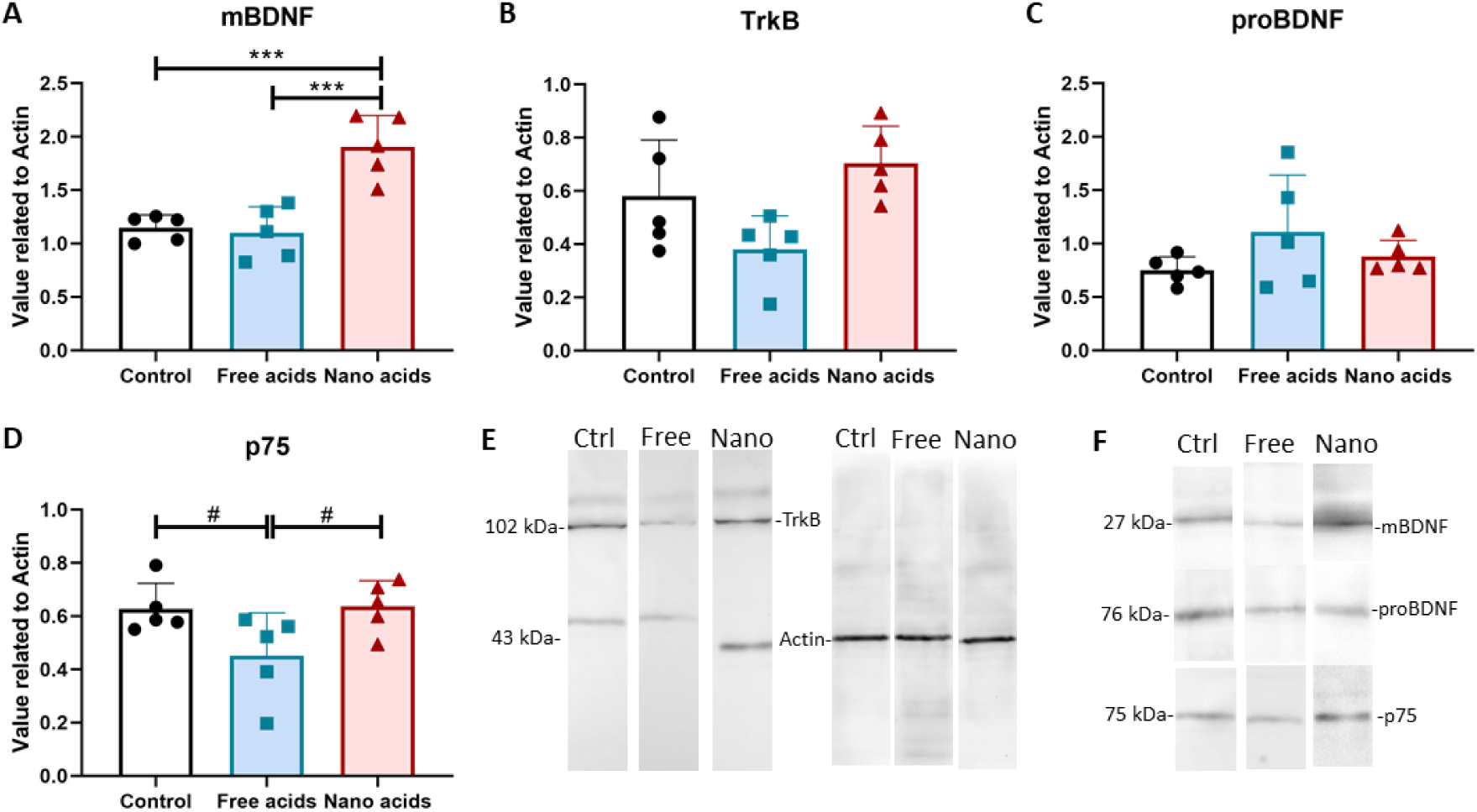
Protein expression of molecules from the BDNF signaling pathway determined by Western Blot and representative membranes. (A)mBDNF. (B)TrkB. (C)proBDNF. (D)p75 for yogurt treated mice after 4 weeks. (E–F) Representative membranes showing signals for TrkB, mBDNF, pro-BDNF, and p75 in Control and fatty acids-treated mice. Data are expressed as mean ± SD, n =5−6/group. # 0.05 < p < 0.10; *p < 0.05; **p < 0.01; ***p < 0.001.

## Discussion

The results presented herein demonstrate that chronic administration of nanoencapsulated PUFAs to adult mice increases the bioavailability of DHA that, in turns, may promote the survival of the newborn neurons in the HC by enhancing the pro-neurogenic elements of the BDNF pathway.

Although there are controversies regarding establishing a recommended dietary intake range for the omega-6/omega-3 ratio (18–21), Zhang et al. (2024), in a UK Biobank large population cohort study, revealed a strong association between the plasma omega-6/omega-3 PUFA ratio and the risk of all-cause mortality, cancer, and cardiovascular disease (22). This supports dietary interventions to increase omega-3 fatty acid levels and maintain a lower omega-6 to omega-3 fatty acid ratio as part of a healthy lifestyle.

In *Wistar* rats fed for 10 days a normocaloric but fat-unbalanced diet rich in saturated fats from butter, supplementation with fish oil (EPA + DHA) or chia oil (ALA) increased circulating levels of EPA and DHA compared to the non-supplemented group. These results indicate that the lipid composition of food rapidly influences the serum fatty acid profile even in a diet rich in saturated fats (14). Similarly, in the present rodent model, although PUFAs quantities ingested by different animals may have varied depending on the amount of cube ingested, EPA showed a trend toward higher serum concentrations in the supplemented groups, and the nanoencapsulated yogurt formulation produced a significant increase in serum DHA levels. Consequently, the group consuming the nanoencapsulated omega-3 yogurt exhibited a higher total n-3 fatty acid content and a lower n-6/n-3 ratio, relative to the other groups (total n-3) and to the control group (n-6/n-3 ratio). Indeed, in vitro studies have already demonstrated increased digestibility and bioaccessibility of n-3 PUFAs when administered in nanoemulsions (7,23). Our finding suggests that nanoencapsulation may protect DHA from degradation during storage and digestion, thereby enhancing its stability and facilitating intestinal absorption, as reflected by increased serum DHA levels. Nanoencapsulation systems might protect PUFAs from gastric degradation and optimize intestinal micellization, significantly increasing their concentration (7). The increased serum omega-3 levels in murine models stem from a tripartite synergy between the PUFAs, the soy lecithin nanoliposomes, and the yogurt matrix. In the acidic environment (pH≈4.5), nanoliposomes provide a robust physical barrier against oxidative and proton-induced degradation (24). This is reinforced by a “protein-lipid corona”, formed via hydrophobic and electrostatic adsorption of caseins and whey proteins, which ensures steric stabilization and prevents coalescence during storage and gastric transit (25). In the gastrointestinal tract, these phosphatidylcholine-rich vesicles function as high-efficiency delivery vehicles. Unlike bulk oils, they facilitate rapid micellization, effectively bridging the unstirred water layer to deliver concentrated PUFAs to the enterocyte apical membrane (26,27). At the cellular level, soy phospholipids upregulate FABP, optimizing intracellular trafficking (28). Furthermore, lecithin provides essential precursors for chylomicron assembly, enhancing fatty acid re-esterification and accelerating lymphatic exocytosis. Consequently, this integrated mechanism—from the protective dairy-protein corona to facilitated transport—underpins the superior systemic bioavailability observed (29).

Positive correlations between plasma and brain levels of DHA were already described (30,31). Therefore, we could assume that the observed higher plasmatic levels of PUFAs may have a correlate in the brain of treated mice. In the present study, the FABP-5, a key regulator of DHA uptake in the brain, showed a trend to increase only in the group that received the nanoencapsulated PUFAs. As this group showed the higher plasmatic levels of DHA, this result could altogether show a better bioavailability and distribution of Omega-3 fatty acids under the new formulation. In an elegant series of studies, Pan et al. demonstrated up-regulated expression of brain microvascular FABP5 protein associated with increased brain DHA levels in adult mice fed with a DHA-enriched diet for 21 days (32). Later they reported that mice knock-out for the Fabp5 gene exhibited impaired performances in memory tasks, associated with a reduction in endogenous brain DHA levels (33). These results, together with ours support the hypothesis that increased plasmatic DHA promotes the expression of FABP-5 in the brain, facilitating in turn the increase of DHA levels in the brain.

DHA has been inversely associated with cognitive decline in clinics (34). Indeed, populations with a high consumption of fatty fish or fatty acid supplements display lower incidence of neurodegenerative diseases compared to those consuming diets poor in these nutrients (35). The process of neurogenesis offers a promising avenue in the search for therapies for aging and neurodegenerative diseases, either by replacing lost neurons or by enhancing brain plasticity. The generation of newborn neurons is well characterized in the adult hippocampus, particularly in the DG, where several factors like neurotrophines, have been shown to facilitate the proliferation and/or survival of newborn neurons (see rev in (36). We were not able to see changes in the proliferation rate of DG neurons in adult mice receiving PUFAs, but mice treated with the nanoencapsulated n-3 Omega fatty acids displayed a trend to an increase in the survival of 4-week old neurons, compared with the control group. The enlarged neurogenesis was accompanied by a significant increase of immature and mature markers of neuronal differentiation like Nestin and Calbindin, respectively. Expression of both markers was also enhanced in mice receiving the free acids, revealing that, globally, the n-3 omega fatty acids appear to act at different stages of the neurogenic process, i.e, differentiation, and survival of newborn neurons. Increased neuron survival has been already shown in a model of strong deficit of PUFA where rats fed a fish-oil deficient diet for three generations were then, supplemented with DHA for 7 weeks (37). Another report from rats fed a DHA and EPA-enriched diet during 4 weeks revealed an increase in hippocampal levels of the BDNF as measured by ELISA (10). Similar results were observed for BDNF mRNA and protein levels in a transgenic mice model of Alzheimer’s disease after 2 months of a DHA-enriched diet (11) and in C57BL/6 mice chronically supplemented with lysophosphatidylcholine-DHA or -EPA (38,39). Our results shed light on these reports describing plastic effects of PUFAs in rodents, as we were able to go deep in the analysis of the BDNF biological pathway. In this study, increased levels of total BDNF mRNA were detected in both groups of mice that received the n-3 omega fatty acids compared to the control groups. However, when the level of proteins were analyzed, mice receiving the free fatty acids showed a sort of activation of the BDNF pro-apoptotic cascade, with a tendency towards higher levels of the p-75 receptor and something similar for pro-BDNF compared to the other experimental groups. In this line, a previous study in a mice model of Alzheimer disease found that the administration of a DHA/EPA-enriched diet for 2 months decreased the expression of pro-apoptosis factors like Bcl2, Caspase 3, and Caspase 9 (11). On the contrary, the proneurogenic elements of this biological pathway, like mBDNF and its receptor TrkB, were significantly increased after chronic administration of nanoencapsulated PUFAs, in synthon with the increased survival of the newborn neurons detected in the DG.

## Conclusions

In this work, nanoencapsulated PUFAs orally administrated to male mice for 4 or 8 weeks showed better bioavailability when compared to administration of free acids, and this potentially due to an increase of the specific transporter, the FABP-5 particularly in the HC. Furthermore, the nanoformulation enhanced the survival of the DG newborn neurons, possibly in response to an improved proneurogenic balance of BDNF actions. These positive effects can be key to facilitate memory processes and also, to prevent cognitive decline that can occur at advanced ages.

## Notes

### Competing Interest Statement

The authors have declared no competing interest.

### Summary of Updates

Figure 1 revised and corrected error, Table 2 revised and corrected error, Discussion updated, References revised and corrected.

### Funding

- INTA, Argentina: PE-088, PD-L04-I121, PE- 119
- ANPCyT, Argentina: PICT 2019-0398
- International Brain Research Organization: IBRO Parenthood 2025
- CONICET: PIP 2021-2023 3123

